# Relationships between puroindoline-prolamin interactions and wheat grain hardness

**DOI:** 10.1101/830265

**Authors:** Nathalie Geneix, Michèle Dalgalarrondo, Caroline Tassy, Isabelle Nadaud, Pierre Barret, Bénédicte Bakan, Khalil Elmorjani, Didier Marion

**Author notes:** Corresponding authors: Didier MARION, Nathalie GENEIX UR1268 Biopolymers Interactions Assemblies, La Géraudière, BP 71627, 44316 Nantes, France.

## Abstract

Grain hardness is an important quality trait of cereal crops. In wheat, it is mainly determined by the *Hardness* locus that harbors genes encoding puroindoline A (PINA) and puroindoline B (PINB). Any deletion or mutation of these genes leading to the absence of PINA or to single amino acid changes in PINB leads to hard endosperms. Although it is generally acknowledged that hardness is controlled by adhesion strength between the protein matrix and starch granules, the physicochemical mechanisms connecting puroindolines and the starch-protein interactions are unknown as of this time. To explore these mechanisms, we focused on PINA. The overexpression in a hard wheat cultivar (cv. Courtot with the *Pina-D1a* and *Pinb-D1d* alleles) decreased grain hardness in a dose-related effect, suggesting an interactive process. When PINA was added to gliadins in solution, large aggregates of up to 13 µm in diameter were formed. Turbidimetry measurements showed that the PINA-gliadin interaction displayed a high cooperativity that increased with a decrease in pH from neutral to acid (pH 4) media, mimicking the pH change during endosperm development. No turbidity was observed in the presence of isolated α– and γ-gliadins, but non-cooperative interactions of PINA with these proteins could be confirmed by surface plasmon resonance. A significant higher interaction of PINA with γ-gliadins than with α–gliadins was observed. Similar binding behavior was observed with a recombinant repeated polypeptide that mimics the repeat domain of gliadins, i.e., (Pro-Gln-Gln-Pro-Tyr)_8_. Taken together, these results suggest that the interaction of PINA with a monomeric gliadin creates a nucleation point leading to the aggregation of other gliadins, a phenomenon that could prevent further interaction of the storage prolamins with starch granules. Consequently, the role of puroindoline-prolamin interactions on grain hardness should be addressed on the basis of previous observations that highlight the similar subcellular routing of storage prolamins and puroindolines.

## Introduction

Grain hardness is a major quality trait that determines the milling properties of wheat and subsequent end-uses of flour. Hardness is genetically controlled and the corresponding major locus (*Ha* locus) is located on the short arm of chromosome 5D, encoding the puroindoline genes [1] Two major proteins are expressed, puroindoline A (PINA) and puroindoline B (PINB). PINA and PINB are small (approximately 13 kDa based on their amino acid sequence) cationic proteins (pI around 10), containing five disulfide bonds and a tryptophan-rich domain (TRD). TRD contains five Trp in PINA and three Trp in PINB [1–3]. Deletion of the *Pina-D1* gene or single mutation of the *Pinb-D1* gene is associated with a hard phenotype, whereas the presence of both *Pina-D1a* and *Pinb-D1a* genes is associated with the soft phenotype [1]. Single mutations can affect the TRD (*Pinb-D1b, Pinb-D1d*) or small hydrophobic domains (*Pinb-D1c*). Null mutations of the *Pinb-D1* gene, which is also associated with a hard phenotype, are due to the insertion of a stop codon in the coding sequence leading to the absence of PINB [1].

Wheat hardness is related to the compactness of the endosperm starch-protein matrix, i.e., to the capacity of the protein matrix to cover most of the starch granule surface and/or to the adhesion strength between starch granules and the protein matrix. The latter is mainly composed of storage proteins, i.e., aggregated monomeric gliadins and polymeric glutenins. The physicochemical mechanisms underlying the relationships between PINs, starch-storage protein interactions and grain hardness are still in debate. Originally, it was observed that the 15 kDa proteins, known as friabilins and mainly composed of puroindolines [4], were absent on the surface of starch granules isolated from hard wheat, whereas they are present on starch granules isolated from soft wheat [5]. It was suggested that friabilins/puroindolines associated with the surface of the starch granules could prevent adhesion with the protein matrix [5]. It was then established that puroindolines were proteins that strongly bind to wheat membrane lipids *in vitro* [6–7]. Consequently, it was suggested that the interactions between lipids and puroindolines could play a major role in grain hardness [1,8]. This hypothesis was strengthened by previous results that showed the similar partition of friabilin and polar lipids at the surface of starch granules in soft and hard wheat [9] and, later, by the lower penetration of mutated PINB into anionic phospholipid monolayers [10].

However, this assertion is challenged by a number of facts. First, during starch extraction, only a small part of friabilin/puroindoline was found on the starch granule surface, whereas puroindolines were mainly recovered in the gluten fraction [11]. Second, high-resolution electron microscopy showed that the majority of PINs were associated with the protein matrix and that during endosperm development, storage proteins and puroindolines follow the same route through the secretory pathway to protein bodies and vacuoles [12–13]. Furthermore, using isogenic Falcon hard and soft lines, it was established that glutenin polymer size is closely related to the presence or absence of PINA [13]. Recently, on the basis of the extractability of prolamin aggregates by SDS and front-face tryptophan fluorescence spectroscopy, it was suggested that the interactions between PINs and gluten proteins could provide an additional mechanistic rationale for the effects of PINs on kernel hardness [14].

Wheat gluten proteins are commonly divided into polymeric glutenins (high and low-molecular types) and monomeric gliadins α/β–, γ– and ω– types). The primary structures of gliadins show a high degree of homology, with a number of amino acid deletions and substitutions. These proteins consist of repetitive sequences rich in glutamine and proline (i.e., QPQPFPQQPYP, QPQQPFP, PQQPFPQQ for α/β–, γ–, and ω-type gliadins, respectively) and associated with a non-repetitive domain stabilized by intramolecular disulphide bonds. Glutenin polymers involved in intermolecular disulfide bonds between the high and low molecular weight subunits and glutenins also contain proline-rich repetitive domains [15]. It is worthwhile recalling here that proline-rich polypeptides are known to interact with tryptophan protein side chains [16–17].

Consequently, there are converging facts concerning the role of protein-protein interactions on kernel hardness. Notably, this was already suggested regarding the aggregative propensity of puroindolines *in vitro* [18–20]. We recently demonstrated that a single mutation in PINB can strongly impact puroindoline self-assembly. Indeed, in aqueous media, mutated PINB proteins assemble in large aggregates, whereas PINA and wild-type PINB form monomers or dimers [20]. Furthermore, even in excess of PINA, PINA and hard-type PINB interact to form large aggregates that could account for the dominance of PINB mutations or PINA deletions that lead to a hard phenotype. With regard to these aggregative properties, the extent of lipid monolayer penetration, when PINA and PINB mixtures are used, is significantly decreased in the presence of mutated PINB than in the presence of wild-type PINB [21]. Consequently, these aggregative properties may also impact the interactions of puroindolines with prolamins on their common route to form the starch-protein matrix during endosperm development.

The objective of this work was therefore to focus on the role of PINA on grain hardness and to explore *in vitro* the protein-protein interactions, i.e., PINA-prolamin interactions, as a potential leading mechanism to control grain hardness. This was first done by measuring grain hardness on selected transgenic wheat lines overexpressing different amounts of PINA and then by characterizing the interactions of PINA with gliadins and their major proteins, i.e., α- and γ-gliadins, coupling turbidimetry, light scattering and surface plasmon resonance.

## Materials and methods

### Materials

All high-quality chemicals were purchased from Sigma-Aldrich and Merck. Isolated wheat proteins, i.e., PINA, total gliadin fraction and α– and γ-gliadins, were purified from the hard wheat *Triticum aestivum* cultivar Recital provided by INRA (Le Rheu, France). PINA was purified and characterized as previously described [20].

### Genetic constructs and production of transgenic wheat lines

The gene specifying the PINA protein was isolated from the Renan wheat variety. This hard variety harbors a wild-type *Pina* gene, *Pina-D1a*. After cloning and sequencing, the *Pina-D1a* gene was inserted in the expression vector between the Bx7 promoter (controlling the expression of a wheat high-molecular-weight glutenin allowing thus both specific and significant levels of expression in the starchy endosperm) and the nopaline synthase (nos) terminator. Once obtained and amplified, this genetic construction was introduced by biolistics into immature embryos of the Courtot cultivar collected 15 days after anthesis according to a previously described procedure [22]. Courtot is a hard cultivar with the *Pina-D1a* and *Pinb-D1d* alleles.

After regeneration and selection, two years were required to obtain transgenic T4 generations, the most genetically stable. Plants were sown in pots and grown in a greenhouse with a natural photoperiod under irrigated conditions. All the subsequent characterizations were done on these transgenic T4 generations. A null-segregant line was used as the control since it was submitted to the same genetic events while having lost the transgene due to genetic segregation.

### Grain hardness

Grain hardness values were determined by two methods – the particle size index (PSI) and near-infrared reflectance spectrometry (NIRS, AACC method, 39-70A, 1999) - from whole meal flour ground in a Cyclotec 14920 mill (Hilleröd, Denmark). For PSI, flour was sieved on a 0.075-mm sieve for 10 minutes using a sieve shaker. The sieve flour was weighed and the PSI value was determined as the percent mass of the flour sample. For hardness measurements, grains from the mid part of the spike were collected from four plants and three-four spikes per plant.

### Protein and lipid contents

Protein content was determined using the Kjeldhal method and lipid contents, i.e. starch and non-starch lipids, were determined as fatty methyl esters by gas chromatography on the flour used for hardness measurements [23]. Puroindoline contents were determined by ELISA according to a previously described procedure [24].

### Extraction and purification of gliadins

The wheat flour (500 g) was defatted with 2.5 L of cold acetone for one hour. After filtration on a Buchner funnel, the slurry was re-extracted with 2 L of methylene chloride overnight at room temperature, followed by filtration. The defatted flour was dried overnight under a fume hood. The defatted flour was stirred for 2 hours at 4°C with 2 L of 100 mM Tris-HCl pH 7.8, 0.3 M NaCl and 5 mM EDTA. The solution was centrifuged at 8,000 rpm for 30 min and the pellet was washed twice with deionized water. After centrifugation, gliadins were extracted by stirring the pellet with 2 L of 70% ethanol for 1.5 hours at 4°C. The solution was centrifuged at 8,000 rpm for 30 min. Gliadins were precipitated overnight at −20°C from the supernatant with five volumes of acetone. The protein pellet was then washed twice with deionized water to remove the residual acetone and freeze-dried. The α- and γ-gliadin fractions were purified from the total gliadin fraction according to the procedure described by Popineau and Pineau [25]. The different fractions were dialyzed twice against water containing 0.01 M acetic acid and freeze-dried. The purity of α- and γ-gliadins was finally controlled by acid- and SDS-PAGE (**Fig. S1**).

### Production of the repetitive domain of wheat gliadins

A synthetic gene encoding eight repeats of the pentapeptide PQQPY was produced and purified as described by [26], except for some modifications for the final step in reversed phase chromatography. Briefly, after cleavage of the thioredoxin-DP-(PQQPY)_8_ fusion protein in 70% formic acid, the mixture was freeze-dried. The lyophilized powder was resuspended in H_2_O containing 0.09% TFA (v/v) and applied to a 6-ml C18-T cartridge (Solid Phase Extraction Strata C18-T, Phenomenex, Torrance, CA, USA) previously equilibrated with solvent A (H_2_O, 0.11% TFA v:v), and the (PQQPY)_8_ fraction was eluted with 40% solvent B (acetonitrile, 0.09% TFA v:v) and analyzed by reversed-phase HPLC, as described for the characterization of gliadin extracts. The purity of the synthetic polypeptide was controlled by mass spectrometry (electrospray ionization source with an ion trap mass spectrometer (ESI-MS); LCQ Advantage Thermo-Finnigan, San Jose, CA, USA), as previously described [27].

The purity of recombinant polypeptide (PQQPY)_8_ was checked by RP-HPLC (**Fig. S2**) and mass spectrometry. Notably, the mass spectrometry (ESI-MS) measurements perfectly matched the expected mass: MW = 4924.5 Da (theoretical mass = 4924.3 Da).

**Particle size analysis of protein aggregates by dynamic light scattering (DLS) and laser light diffraction**.

**The particle size of protein aggregates was measured using a Malvern Zetasizer Nano ZS90 for DLS and a Mastersizer S Instruments (Malvern Instruments, UK) for laser light diffraction. DLS measures hydrodynamic diameters of particle size below 100 nm [20] and the laser light diffraction measures aggregates ranging from hundreds of nanometers up to several millimeters in size. The angular scattering intensity data is then analyzed to calculate the size of the particles using the Mie theory of light scattering. The analysis requires the relative refractive index of the continuous (water) and the dispersed phase: 1.33 and 1.52, respectively. The particle size is reported as a volume equivalent sphere diameter. Dilute solutions of PINA and gliadins (0.5 mg.mL^−1^) in 20 mM Tris-HCl, pH 7.2, were measured in a 1-cm path-length spectroscopic plastic cell (BRAND, France) at 25°C and for 30 s for DLS. By using laser light diffraction, 1 mL of concentrated sample with a PINA/gliadin ratio of 1:10 (w:w) was diluted with 15 mL deionized water in the sample dispersion curve under stirring.Turbidity measurements.**

The concentration of PINA was calculated by measuring optical density at 280 nm with a UV-spectrophotometer (Perkin Elmer Lambda 2) and by using an extinction coefficient of PINA (31.10 M^−1^ cm^−1^ or 2.376 g^−1^ L.cm^−1^). The gliadin solution concentration was adjusted as a function of optical density at 280 nm of standard solution of gliadin (1 mg.mL^−1^, DO~0.4). Gliadins were solubilized in 70% ethanol to achieve a final concentration of 20 mg.mL^−1^ and were then diluted at 2 mg.mL^−1^ in 20 mM Tris-HCl at pH 7.2, 20 mM malate at pH 5, or 20 mM citrate at pH 5 buffer solutions. Purified PINA was solubilized at 1 mg.mL^−1^ in these different buffers. Increased quantities of PINA were added, and after stirring for 30 min at room temperature, the optical density at 540 nm of 0.1 mL mixture solution was measured in a 96-well microplate reader Epoch (BioteK® Instruments, USA).

### Surface plasmon resonance (SPR) measurements

SPR measurements were performed on a SPR X100 instrument (GE-Healthcare Biacore) with an automatic flow injection system. SPR buffers, regeneration solutions and Sensor Chips coated with carboxylated dextran (CM5) were purchased from GE Healthcare (France). A quantity of 250 µL of gliadins (20 µg.mL^−1^) in 10 mM acetate buffer (pH 4.7) were covalently bound to the first channel of the chip surface by esterification using carbodiimide crosslinker chemistry, i.e., N-(3-Dimethylaminopropyl)-N′-ethylcarbodiimide hydrochloride/N-hydroxysuccinimide [28]. The other control channel received no protein. After immobilization, residual free sites were blocked by injection of 70 µL of 1 M ethanolamine at pH 8.5 for 6 min. First, initial buffer flow gave a reference baseline. Second, the interaction between PINA and gliadins (α- or γ-gliadin or recombinant polypeptide (PQQPY)_8_) was studied at 25°C by flowing various concentrations of PINA over the immobilized gliadin sensor surface for 1 min at a flow rate of 10 µL.min^−1^ in Hepes buffer, pH 7.4 (10 mM Hepes, 150 mM NaCl, 3.4 mM EDTA, and 0.005% P20 surfactant). The sensor surface was regenerated between sample injections by flowing 50 mM NaOH for 0.5 min at a flow rate of 10 μL min^−1^. The formation of a PINA/gliadin surface-bound complex was detected by refractive index and changed the resonance signal (expressed in resonance unit RU) as a function of time (sensorgrams). After 120 s, a steady-state plateau was established where association and dissociation of protein-protein complexes occur at an equal rate. Third, a free-PINA buffer flow was repeated and the dissociation of the complex was monitored as a function of time. The sensorgrams for the different PINA concentrations were overlaid and aligned to the baseline determined in the absence of PINA (Fig. S3).

Using raw data, the binding kinetic parameters of PINA to gliadins were determined: k_a_, k_d_, K_D_=k_d_/k_a_, i.e., association rate, dissociation rate and equilibrium dissociation constant, respectively. They were determined by fitting the experimental data with the 1:1 Langmuir model (BIA-evaluation software 4.1, GE Healthcare). The binding constants were calculated by two methods: affinity analysis (steady-state fit) and association/dissociation analysis (kinetic fit) [29].

## Results

### Dose-response effect of the over-expression of PINA on wheat hardness

PINA was overexpressed in the Courtot genetic background. This cultivar was chosen because its endosperm is particularly "hard”. Indeed, it bears a mutant version of the *pinb* gene (*Pinb-D1d*), where a mutation leads to the substitution of a tryptophan side chain by an arginine at position 44, i.e., in the TRD that determines the physicochemical properties of PIN [1]. Five transgenic lines of the 4^th^ generation were selected that express PINA at different levels when compared to their respective null-segregant. This set of transformants offers the opportunity to investigate the dose effect of PINA on grain hardness.

First of all, it is essential to check the structure of overexpressed puroindolines to determine their functionality on grain texture. PINs are notably known to be the targets of a series of successive post-translational maturations consisting of cleavages at both their N- and C-termini [2,20]. In previous studies, the overexpression of PINs was checked by using SDS-PAGE, eventually coupled with Western blotting [30]. Such techniques are of course not sufficient to characterize the structure of the overexpressed protein. We therefore checked whether this post-translational processing is affected by the expression of the transgene. Indeed, the maturation of the protein was ensured, as previously reported in different wheat cultivars [2, 20], except for some slight differences in the maturation of the C-termini (Fig. S4). Compared to the null-segregant, the ratio of the least truncated vs. more truncated forms of the PINA extracted from the transgenic lines is significantly higher (Fig. S4b). Similar differences in the maturation of PINA between soft and hard varieties have been previously observed [20]. This suggests differences in the regulation of the machinery involved in the post-translational processing of puroindolines and could also be related to the higher expression of PINA in very soft varieties when compared to very hard ones [8]. These changes should not impact the properties of overexpressed PINA, as previously observed when comparing the formation of heterodimers with PINA extracted from soft and hard wheat [20] and actually these modifications are not significantly correlated to the amount of PINA overexpression (Fig. S4b).

Grain hardness measured by both particle size index (PSI) and NIRS was significantly reduced in the lines overexpressing the *Pina-D1a* allele compared to the corresponding null-segregant (Table 1). In the present work, we showed that hardness levels decreased almost proportionally to the expression levels of the PINA protein with a significant correlation factor: an R2 of 0.94 for the PSI and an R2 of 0.7 for the NIRS measurement of grain hardness (Fig. 1). Furthermore, we also showed that the expression of the PINB-D1d is not impacted by the overexpression of PINA. Finally, the impact of PINA overexpression on proteins and lipids was also investigated since PINA interacts with lipids *in vitro* [7,18, 21] and *de facto* with storage proteins *in planta*, where they follow the same intracellular routing pathway [13]. The protein and lipid contents of the five lines are not significantly different from the null-segregrant lines (Table 1). This impact of the overexpression of PINA on hardness was observed for the expression of recombinant PINA in other genetic hard wheat backgrounds [30]. The data indicate that PINA and PINB interact to form friabilin and together affect wheat grain texture. By studying the PINA-PINB interactions, i.e., the formation of heteromeric aggregates, we previously showed that only the hard versions of PINB interact with PINA, a phenomenon that could in part limit the pro-softness properties of PINA [20].

**Table 1.** Hardness and biochemical composition of transgenic lines overexpressing PINA. Grain hardness was determined by particle size index (PSI) and near-infrared reflectance spectrometry (NIRS). The PSI reported is the percentage of flour recovered that is smaller than 75 µm. PSI decreases with the hardness of wheat. The NIRS value increases with the increase in particle size and the particle size of wheat flour increases with hardness.

| Lines | PINA<br>mg % | PINB<br>mg % | PSI<br>(%) | NIRS | Protein<br>(%) | Non-starch<br>lipids % | Starch-<br>lipids % |
| --- | --- | --- | --- | --- | --- | --- | --- |
| A5NS* | 25 ± 4 | 11 ± 1 | 41 | 78 | 17.5 | 1.6 | 0.37 |
| A1 | 70 ± 8 | 15 ± 3 | 45 | 60 | 18.1 | 1.6 | 0.36 |
| A3 | 127 ± 23 | 19 ± 2 | 46 | 56 | 17.4 | 1.7 | 0.37 |
| A2 | 155 ± 10 | 20 ± 6 | 49 | 66 | 18.9 | 1.7 | 0.35 |
| A6 | 163 ± 21 | 22 ± 2 | 50 | 52 | 17.9 | 1.6 | 0.37 |
| A4 | 214 ± 12 | 15 ± 6 | 51 | 42 | 17.1 | 1.7 | 0.36 |

\*(NS) null-segregant. Values are means ± SD (standard deviation, N=4).

**Figure 1.**
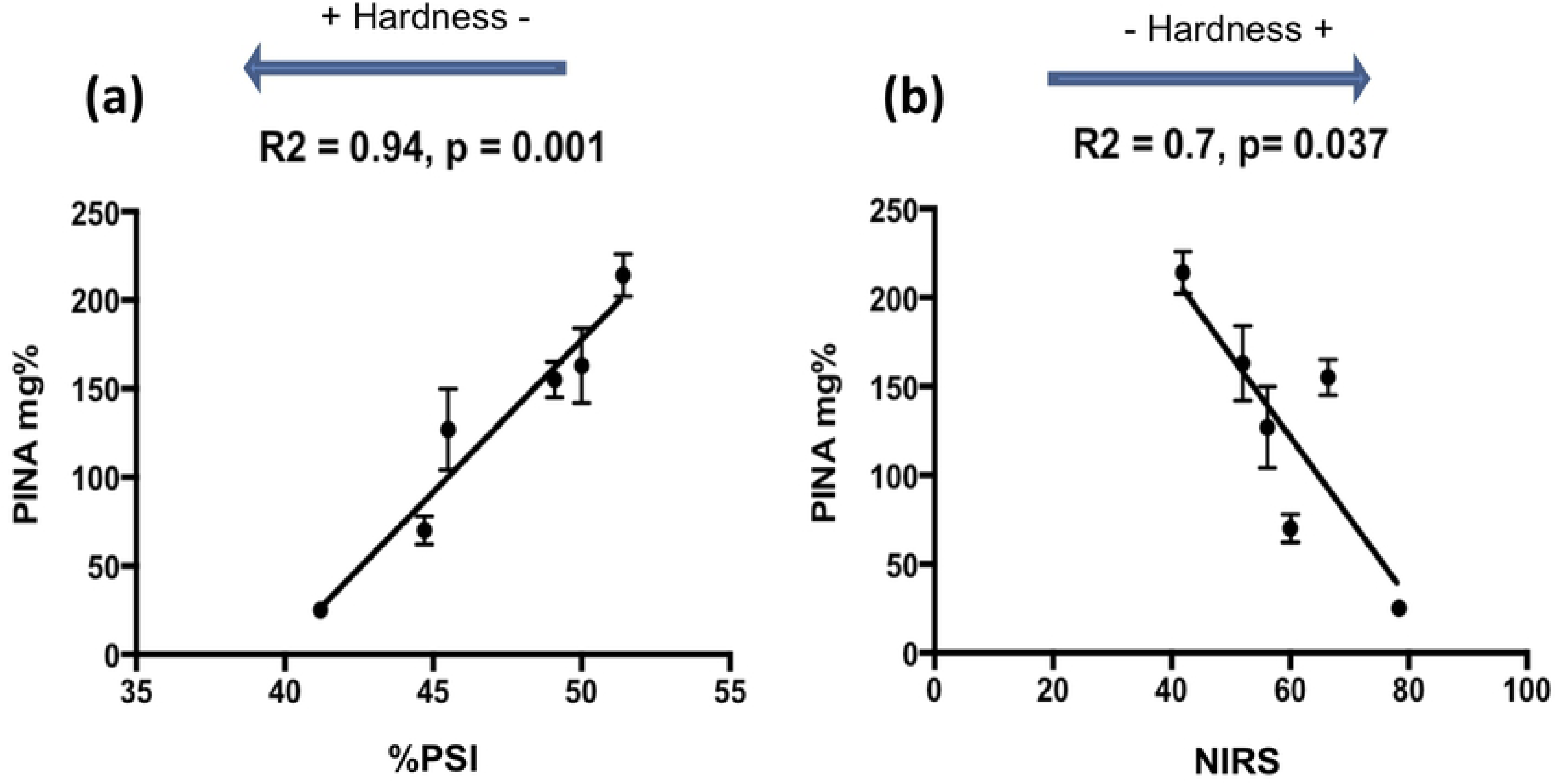
Grain hardness. Correlations between PINA contents and hardness measured through PSI (a) and NIRS (b).

Accordingly, the dose-related effect of PINA on grain hardness suggests an interactive process. Since the puroindolines have been shown to be co-localized in the protein bodies of developing endosperms and mainly in the protein matrix at the end of wheat endosperm development [12–13], we decided to further pinpoint the molecular basis between PIN and endosperm texture through the characterization *in vitro* of the interaction of PIN with gliadins, one of the major protein families of wheat storage proteins.

### PINA impacts the aggregation of gliadins

Gliadins, in contrast with glutenins, are monomeric proteins soluble in hydroalcoholic or acidic solutions and were therefore chosen to explore PINA-prolamin interactions. We first checked the interactions of PINA with the total gliadins extracted from Recital, a hard wheat cultivar. The gliadin fraction is composed of ω, α, β and γ-gliadins, as shown by acid-PAGE, SDS-PAGE and RP-HPLC (**Figure. S1**).

Interactions were assessed by light scattering and turbidity. Indeed, immediately after gliadin solubilization (2 mg.mL^−1^), turbid solutions were observed in the absence and in the presence of 0.2 mg.mL^−1^ of PINA (Figure. 2). This turbidity persists 30 min later for the PINA-gliadin mixture, whereas it partially disappears for the gliadin mixture alone. After 30 min, the PINA solution displays a single particle population with a mean diameter of 8 nm. Immediately after dilution, gliadins form aggregates with a monomodal Gaussian distribution by volume and a mean diameter of around 6 µm. However, the 6-µm gliadin particles dissociate into smaller aggregates, i.e., 6 nm diameter, after 30 min. In the case of a PINA/gliadin (1:10, w:w) mixture, a single population of large particles is obtained with a diameter of around 1 µm immediately after dilution and 13 µm after 30 min of mixing. No further modification of particle size could be observed over a longer time, suggesting that at these relatively low concentrations, gliadin aggregation is certainly a dynamic process controlled by structural rearrangements.

**Figure 2.**
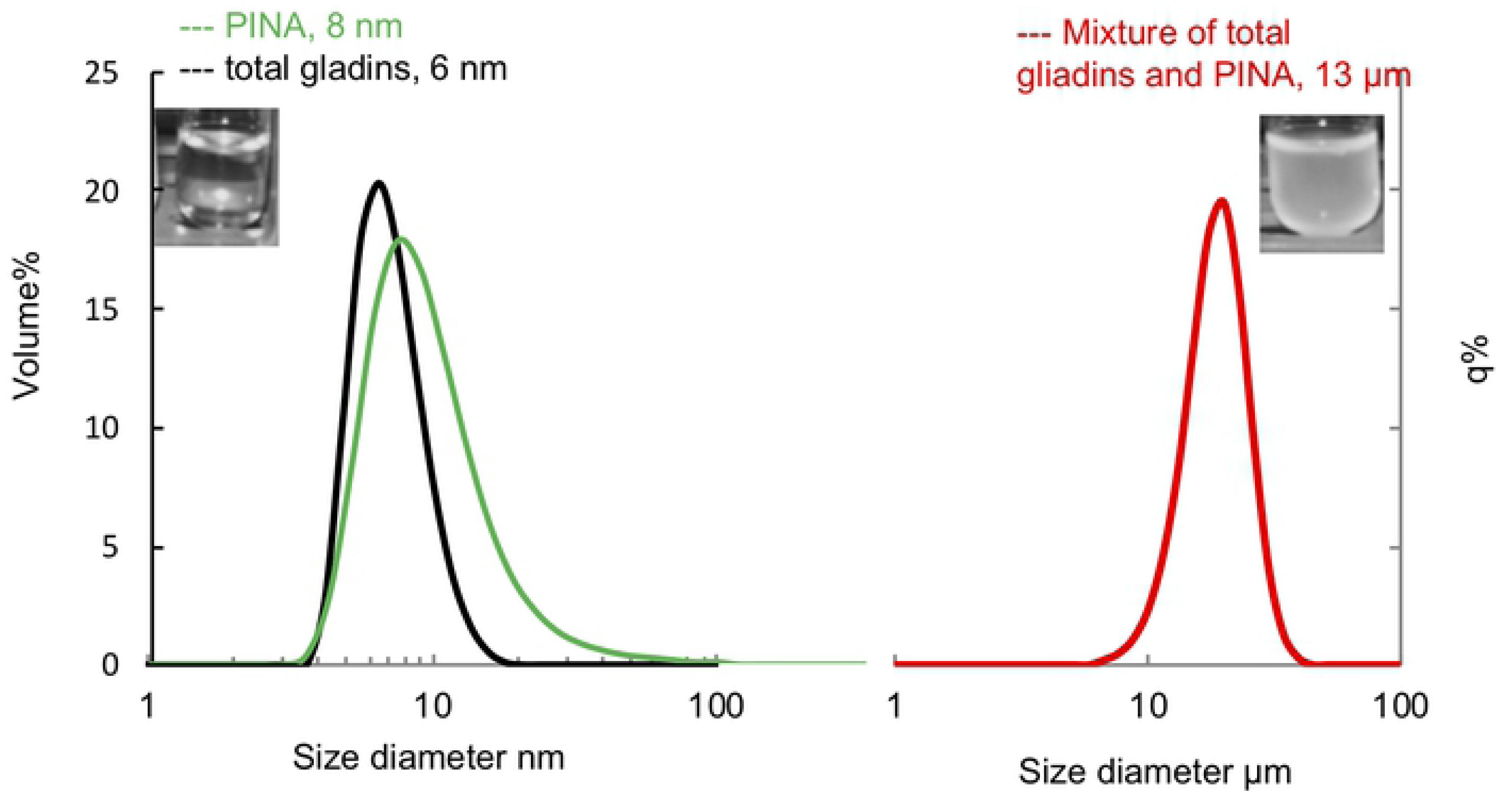
Size distribution of protein aggregates in solution after 30 min of mixing. Measurements of between 1 and 100 nm were carried out by dynamic light scattering for PINA particles (green line), and gliadin particles (black line). Measurements of between 1 to 100 μm were carried out by laser light diffraction for the PINA/gliadin mixture (red line). Inserts: pictures of turbidity solutions of PINA with total gliadins.

To check the dose-related effect of PINA on gliadin aggregation, turbidity measurements were monitored at a constant concentration of gliadins (2 mg.mL^−1^) while increasing concentrations of PINA. In addition, the biological context was considered, especially the endosperm pH that decreases during endosperm development, first slowly up to 30 DPA, and then rapidly to reach about 4.5 at the end of this process (50 DPA). Acidification correlates very well with the accumulation of organic acids, i.e., malate, citrate, lactate [31]. Accordingly, gliadin/PINA interactions were also monitored either in Tris buffer, pH 7.2, in 20 mM malate at pH 5 or in 20 mM citrate at pH 5. As shown in Fig. 3, turbidity increases following the addition of PINA to reach a plateau at a PINA/gliadin mass ratio of 1.5:100 (w:w) at pH 7.2 and 0.8:100 (w/w) at pH 5. The slight lag concentration effect when PINA is added to the gliadin solution suggests that the aggregation process is highly cooperative. Indeed, curve fitting using the Hill ligand-binding model gives rise to a Hill coefficient of 1.9 at a neutral pH and of about 3.5 at an acidic pH (Fig. 3). This high cooperativity is probably associated with a nucleation effect, i.e., the formation of the first PINA-gliadin complexes that promote further aggregation of other gliadins. These titration curves were not impacted by the presence of salt (data not shown), suggesting that the aggregation process was not driven by electrostatic interactions but only by the combined action of organic acids and pH.

**Figure 3.**
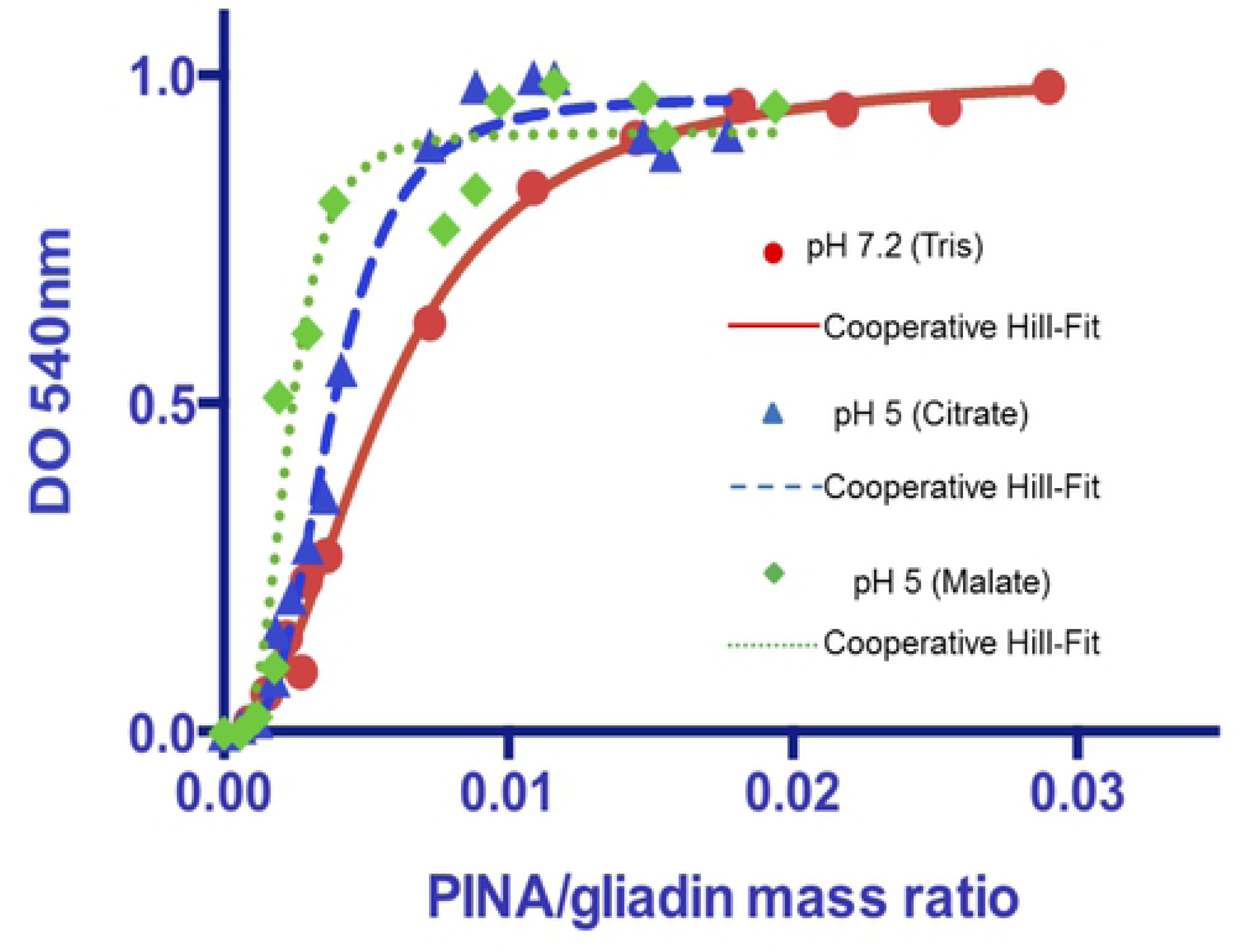
Turbidimetry measurement (optical density at 540 nm) after 30 min mixing of gliadins with increased PINA concentration. Curve fits for PINA-total gliadins give a Hill coefficient of 1.9 (Tris pH 7.2), 3.3 (citrate pH 5) and 3.7 (malate pH 5). A Hill coefficient > 1 indicates a cooperative binding. K_D_ was measured: 1 μM (pH 7.2), 0.6 μM (pH 5) and 0.4 μM (pH 5).

### Selectivity and role of the gliadin repetitive domain in PINA-gliadin interactions

Gliadins refer to four protein families, i.e., α, β, γ and ω gliadins, owing to their electrophoretic mobility in acid-PAGE and their amino acid sequences [15] (**Fig. S2**). We showed that PINA interacts with total gliadins to form large aggregates. Since α– and γ-gliadins are the major proteins in the gliadin fraction, they were chosen to probe their interaction with PINA. In contrast with total gliadins, adding PINA to α– or γ-gliadin fractions led only to a slight increase in turbidity (data not shown). This suggests that PINA does not impact the aggregative properties of α– and γ-gliadins, or induces a slight increase of the size of gliadin aggregates, undetectable by turbidity measurements. We therefore checked whether α– and γ-gliadins interact *in vitro* with PINA by using SPR. Figure 4 shows sensorgrams obtained after having increased PINA concentrations (0.3 µM to 5 µM) on α-gliadin- or γ-gliadin-coated chips. Sensorgrams demonstrate that PINA interacts with immobilized α-gliadin or γ-gliadin with a dose-dependent effect and very fast association/dissociation kinetics, with equilibrium binding, i.e., steady-state binding, achieved during the 120-s association phase. PINA/gliadin binding reaches a steady-state plateau during the injection period; the responses are flat and rapidly return to the baseline. At equilibrium and for the same concentration of PINA (i.e., at PINA=2.5 µM), the RU signal is higher for α-gliadin (160 RU) than for γ-gliadin (34 RU). The association rate (Ka) is significantly lower for α-gliadin than for γ-gliadin but the dissociation one (Kd)is in the same order of magnitude. These results suggest that PINA has a higher affinity for γ-gliadin than for α-gliadins.

**Figure 4.**
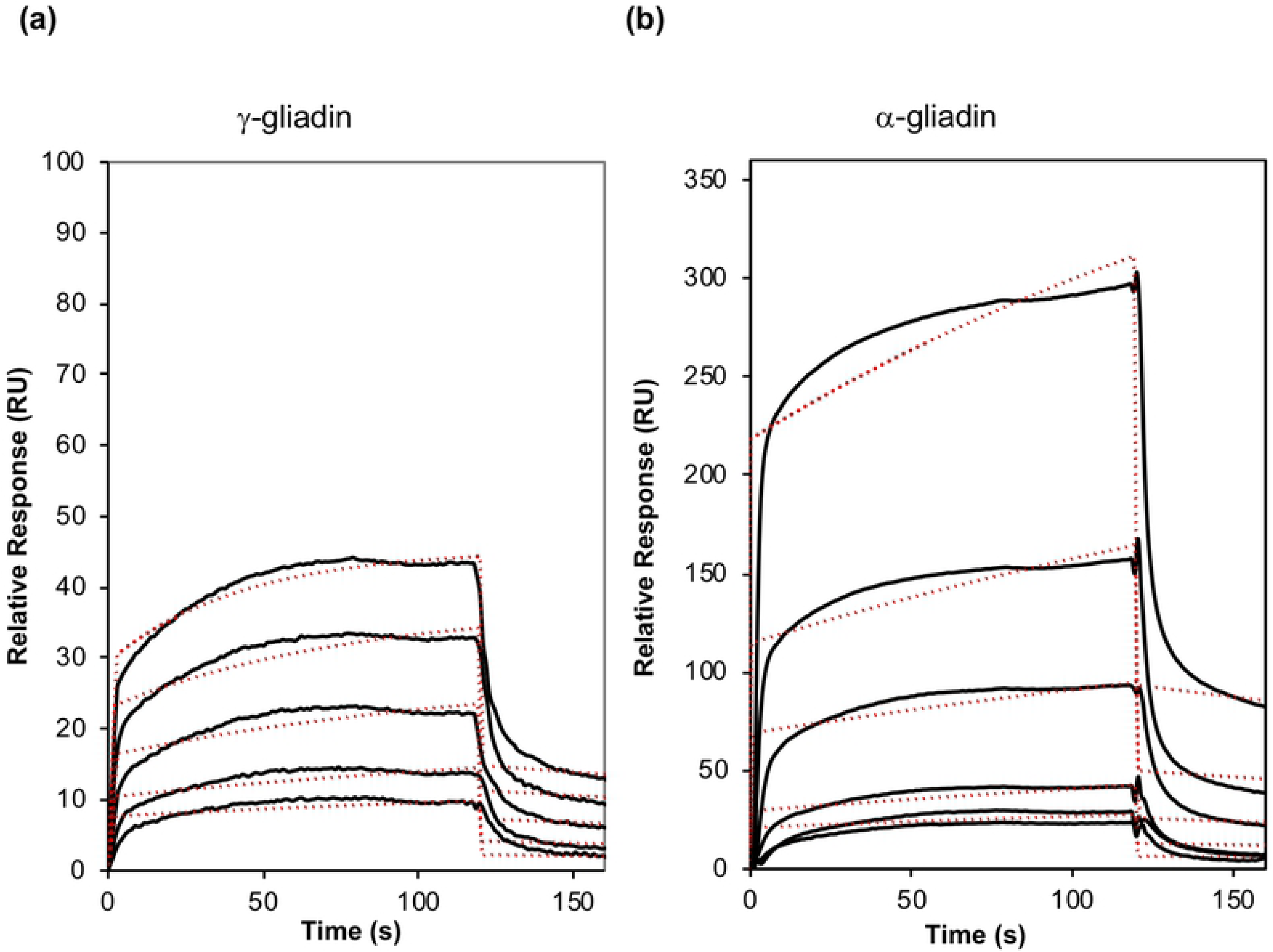
Binding analysis. Sensorgrams for PINA binding to (a) γ-gliadin and (b) α-gliadin. The flow rate was 10 μL.min^−1^, with contact times of 120 s and dissociation times of 600 s, in HBS-EP running buffer. Recorded sensorgrams (black lines) for the different PINA concentrations were overlaid and aligned with the baseline determined in the absence of PINA. Fitting curves are shown in dashed red lines. The PINA concentrations from bottom to top are 0.31, 0.62, 1.25, 2.5 and 5 μM.

The structure of gliadins is characterized by the presence of N-terminal domains composed of repeated peptide motives and the recombinant polypeptide (PQQPY)_8_ mimics the consensus repetitive domain of gliadins. Injection of 2 µM to 20 µM of PINA over the immobilized (PQQPY)_8_ polypeptide (Fig. 5) confirms the interaction of PINA with the repeat consensus sequence. The sensorgram for 2.5 µM of PINA displays a signal intensity that is greater than that of α-gliadin (RU= 300), but return to the baseline is slower for the (PQQPY)_8_ polypeptide.

**Figure 5.**
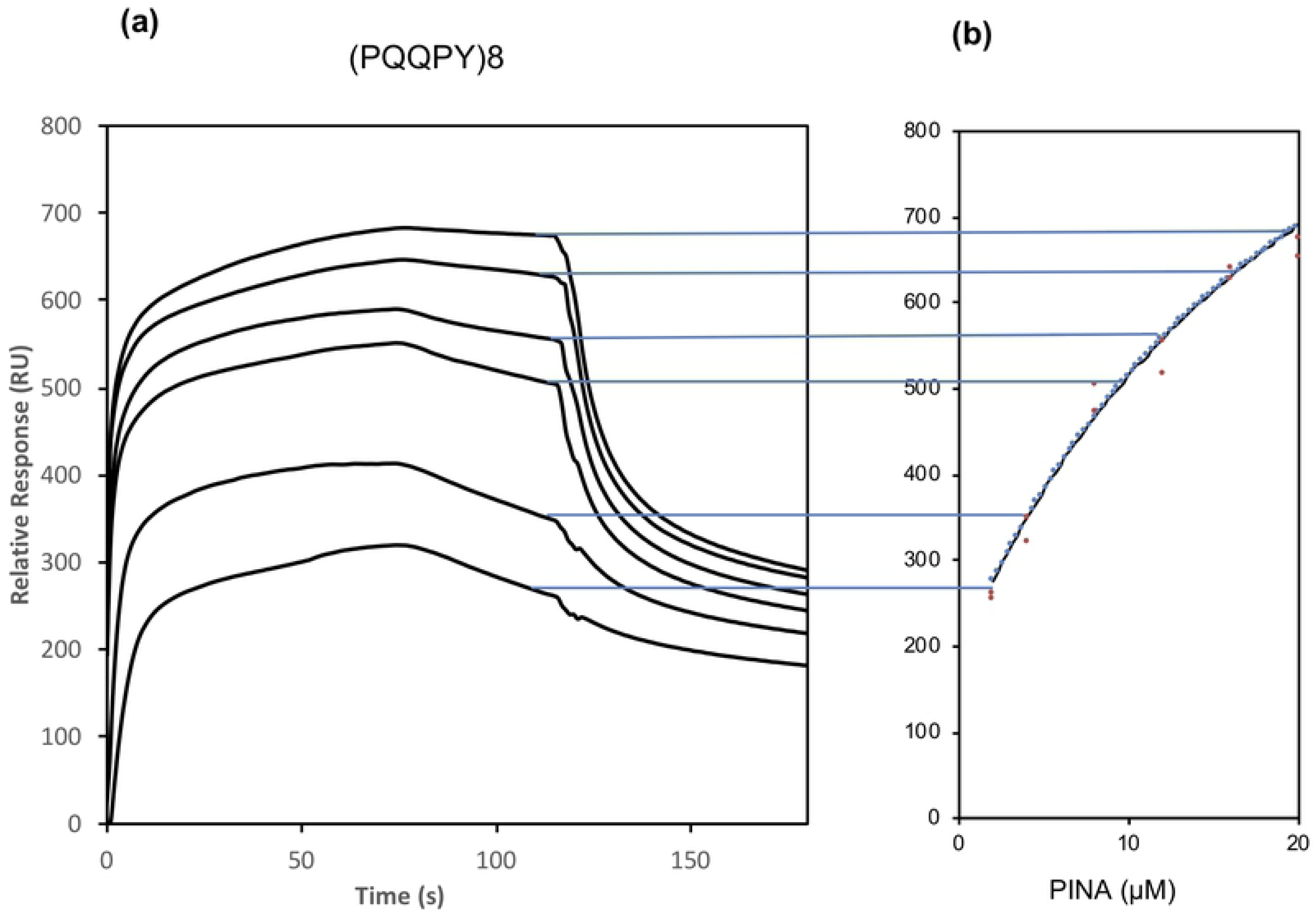
Kinetic analysis. (a) Sensorgrams for PINA binding to (PQQPY)_8_ polypeptide. The PINA concentrations from bottom to top are 2, 4, 8, 12, 16 and 20 μM. (b) Fitting for a steady-state binding Langmuir 1:1 model to estimate the equilibrium dissociation constant K_D_ for PINA binding to the (PQQPY)_8_ polypeptide.

A global kinetic fit with a Langmuir 1:1 binding model was applied for the sensorgrams to determine the binding kinetics, k_a_, k_d_ rate constants and K_D_ (equilibrium dissociation constant, K_D_=k_d_/k_a_) (Figs. 4 and 5, Table 2). The data over a selected period in the steady-state region of each sensorgram (20 s before the end of injection) can be plotted as a function of protein concentration in a flow solution for binding PINA/(PQQPY)8 (Fig. 5). The Langmuir 1:1 binding model was used and can only be fitted on an interaction with a 1:1 stoichiometric ratio. K_D_ with a steady-state fit was also calculated (Table 2).

**Table 2.** Kinetic and affinity (steady state) constants determined by SPR.

| | $k_a$<br>($\times 10^{-3} \text{ M}^{-1}\text{s}^{-1}$) | $k_d$<br>($\times 10^3 \text{ s}^{-1}$) | $K_D$ ($\mu\text{M}$)<br>Kinetic fit | Steady-state fitting |
| --- | --- | --- | --- | --- |
| γ-gliadin | $3.6 \pm 0.9$ | $1.85 \pm 0.5$ | $0.63 \pm 0.1$ | nd* |
| α-gliadin | $0.45 \pm 0.04$ | $2.55 \pm 0.7$ | $4.45 \pm 0.07$ | nd* |
| (PQQPY)<sub>8</sub> | $5.0 \pm 0.03$ | $1.50 \pm 0.1$ | $0.29 \pm 0.005$ | $7.22 \pm 2.8$ |

\*not determined

K_D_ of (PQQPY)_8_ for PINA (0.29 µM) is on the same order of magnitude as γ-gliadin for PINA (0.63 µM) but is more than 15 times lower than that of α-gliadin for PINA (4.45 µM). The dissociation constant K_D_ with the steady-state model (at equilibrium) was calculated for PINA with (PQQPY)_8._ KD (7.22 µM) results are higher than the results determined by kinetic rate constants (KD=kd/ka=0.29 µM). These differences in K_D_ using the steady-state method can be attributed to an insufficient concentration of PINA to saturate the binding sites or to an inadequate binding model to perfectly fit the experimental data. Regardless of the binding model, SPR experiments clearly revealed that PINA interacts with gliadins and especially with their repetitive domain.

## Discussion

PINA was chosen in this work because its absence led to hard kernels and its content is lower in hard wheat bearing a mutation in PINB, as observed here with the Courtot cultivar. Furthermore, *in vitro*, hard variants of PINB form large aggregates that, interestingly enough, can trap PINA and could limit the impact of PINA on hardness [20]. Beyond the confirmation that the overexpression of PINA in hard wheat increases kernel softness, this work clearly shows a dose-related effect of PINA on grain texture. Actually, the reduction of grain hardness is not that high - a maximum decrease of around 30% - as observed by Hogg et al. [30] for the overexpression of PINA in a hard wheat bearing the *pinb-D1b* and *pina-D1a* genes. Indeed, these authors observed a higher impact of soft PINB, i.e., PINB-D1a, and of the association of soft PINA and PINB to significantly decrease the hardness of a hard cultivar [30]. In any case, the dose-related effect of PINA on grain hardness could probably be observed thanks to an ELISA using specific monoclonal antibodies, which ensures the absolute value of PINA and PINB contents. In addition to screening and the selection of transgenic lines with different expression levels of PINA, this ELISA showed that the overexpression of PINA did not impact the expression of the PINB-D1d variant that is characteristic of the hard Courtot cultivar. Therefore, in our experiments, hardness changes are due only to the overexpressed PINA, a result strengthened by the absence of modifications in the contents of the major putative molecular targets of puroindolines, i.e., lipids and proteins, which control the interactions of the protein matrix with starch granules. Regarding the common intracellular routing of puroindolines and storage proteins as well as the aggregative properties of these proteins *in vitro* [18–20], we explored the possibility of PINA-prolamin interactions to account for the dose-related effect of PINA on hardness.

We focused on the interactions of PINA with gliadins since these storage proteins can be easily solubilized in hydroalcoholic or acidic solutions, unlike the insoluble glutenin polymers that impair *in vitro* studies. Our data showed that the presence of PINA strongly impacts the aggregation behavior of gliadins in solution. Using SPR, we demonstrated that PINA can interact with the two major gliadin families, i.e., α-gliadins and γ-gliadins. In addition, we showed that PINA also interacts with a polypeptide that mimics a repeated peptide consensus sequence (PQQPY)_8_ of gliadins. This latter observation suggests that the repetitive domain of these storage protein families is involved in the PINA-gliadin interaction. This result is strengthened by the fact that tryptophan is often involved in the interaction of proteins with proline-rich domains [32–33]. In this regard, the significant difference in the affinity of PINA for α- and γ-gliadins could be due to differences in the consensus repeat peptides in these gliadin families [34]. Finally, the higher affinity of PINA for γ-gliadins could account for their partition with PINs in the detergent-rich phase when applying TX114 phase partitioning for their purification [35].

The most outstanding result of this study was when PINA comes into contact with total gliadins, i.e., with the natural composition of the different gliadin families. Gliadins form nanosized particles in dilute solution and, in the presence of PINA, the particle size increased by a factor of 10 with the formation of aggregates of about 13 µm in size. This behavior allowed turbidimetric measurements to determine the steady-state binding of PINA to gliadins. The corresponding binding curves highlighted a highly cooperative phenomenon. This suggests that initiation of a PINA-gliadin complex could then facilitate the aggregation of the other gliadins. Our present results on α- and –γ gliadins show that this initial nucleation could involve these gliadins and, in particular, their repetitive domain. This cooperativity is more pronounced at acidic pH in the presence of malate and citrate than at neutral pH. Previous studies on barley endosperm have shown an accumulation of organic acids during development, leading to progressive acidification of the endosperm [31]. Interestingly, this cooperative aggregation needs to maintain the protein diversity, i.e., the natural mixture of α-,β-,γ-and ω-gliadins. Indeed, we could not observe any changes in turbidity when PINA was added to individual α- and γ-gliadin solutions or any cooperative binding in SPR experiments with these prolamins. This cooperativity should allow the amplification of prolamin aggregation during endosperm development, thus favoring protein-protein associations during desiccation to the detriment of starch-protein associations. This hypothetical cooperative protein-protein interaction mechanism can also account for the strong effect of minor proteins on the hardness phenotype.

The protein-protein interaction data obtained *in vitro* should be analyzed in view of the observation made in this work that the overexpression of PINA in a genetic background of "hard" wheat, i.e., cv. Courtot, leads to a decrease of grain hardness with a dose-related effect and with previous observations showing that the hard variant of PINB can trap PINA in heteromeric PINA-PINB complexes. The formation of these heteromeric complexes could in part explain the limited effect of PINA on grain hardness through a competition between PIN-PIN and PIN-gliadin interactions. In other words, the excess of free PINA, i.e., not associated with PINB-D1d aggregates, increases with the increase in the overexpressed PINA level and is available for the expression of a softer phenotype. Furthermore, the limited impact of PINA overexpression on hardness could be exemplified by the high protein contents of our transgenic lines related to greenhouse growing conditions. It is in fact known that hardness increases with the increase in protein content [36]. High protein content could limit the extent of cooperative protein aggregation, leaving an excess of storage proteins that interact with the starch granule surface.

The predominant hypothesis suggests that PINs would be adsorbed on the surface of the starch grains, preventing adhesion between the starch grains and the protein matrix. Due to the presence of lipids on the surface of starch grains (surface starch lipids) and the interactions between PINs and lipids observed *in vitro*, it has been suggested that these interactions between PINs and lipids would prevent the adhesion of the protein matrix to starch granules. This hypothetical mechanism could not be completely ruled out since the affinities of PINA for wheat lipids and for gliadins are on the same order of magnitude, i.e., KD in the micromolar range [6]. The spatial distribution of lipids and proteins is not homogeneous and undergoes changes during endosperm development. However, regarding the routing of PINs from the endoplasmic reticulum to vacuoles where the storage proteins accumulate at the same time, i.e., until 30 DPA [13], PINA and prolamins remain in close proximity, whereas the contact of PINA and lipids should occur only on desiccation when the membranes of subcellular organelles collapse and accumulate at the interface of the starch-protein matrix [37–38]. Furthermore, the impact of PINA on prolamin aggregation can occur during endosperm development since these interactions occur regardless of the pH, from neutral to acidic conditions. It is also worthy to note that, besides the expression level of puroindolines, those of storage proteins, especially gliadins, are different between hard and soft cultivars [39].

It will also be important to further consider the role of the soft PINB variant in these protein-protein and protein-lipid interactions. Although a previous study did not show any physical interactions between PINA and the soft PINB variant, it was clearly demonstrated that only the overexpression of both PINA and the soft PINB variant are necessary to transform a hard wheat into a very soft cultivar [30]. With regard to our results, this synergy could be due to greater interaction of PINB with prolamins or with lipids covering the surface of starch granules. Indeed, we previously observed a higher affinity of PINB than PINA with wheat polar lipids [6].

## Conclusion

PINA impacts wheat grain texture as demonstrated by the significant correlation that could be established between PINA levels and hardness. By combining turbidimetry and light scattering, we clearly demonstrated that this phenomenon could involve protein/protein interactions. PINA can interact with gliadins and impact their aggregative properties. The high cooperativity of this interaction induces the aggregation of other gliadins to the nucleating heteromeric PINA-gliadin complex. This fact can explain why a small increase of PINA in hard wheat drastically impacts grain hardness with a dose-response effect. Due to gliadin-glutenin associations in the protein matrix, this cooperative phenomenon could be extended to glutenins.

The specificity of these interactions, i.e., PINA vs. gliadin type (α, β and γ-gliadins) and PINA vs. repeat or non-repeat domains, deserves to be further tackled in order to further clarify the role of protein-protein interactions, in particular, the role of PINB and other storage proteins as well, i.e., glutenins, on the formation of the starch protein-matrix *in planta*.

## Abbreviations

DLS: dynamic light scattering
HEPES: 4-(2-hydroxyethyl)-1-piperazineethanesulfonic acid
HPLC: high-performance liquid chromatography
IEC: ion-exchange chromatography
NIRS: near-infrared reflectance spectrometry
PAGE: polyacrylamide gel electrophoresis
PINA: puroindoline A
PINB: puroindoline B
PINs: puroindoline A and puroindoline B
PSI: particle size index
SDS: sodium dodecyl sulfate
SPR: surface plasmon resonance
TRD: tryptophan-rich domain
Trp: tryptophan
WT: wild type

## Aknowledgements

We are grateful to Marielle Merlino and André Lelion for their technical assistance in the initial genotyping of transgenic lines and the purification of puroindolines, respectively. We also wish to thank Gérard Branlard for his commitment and enthusiasm since the beginning of this work. A part of this work was supported by a grant from the Pays de la Loire Regional Council. We also thank Gail Wagman for critical reading and editing of the manuscript.

## Supporting information

**S1 Figure. Characterization of total gliadins, α- and γ-gliadins.** (a) acid-PAGE (b) SDS-PAGE, and (c) C18-PFP RP-HPLC.

**S2 Figure. Purification of the repetitive domain of α-gliadin.** (a) Coomassie blue stained SDS–PAGE of (1) BLR strain carrying pETb (PQQPY)_8_ plasmid, (2) non-adsorbed affinity column, (3) elution of TRX-(PQQPY)_8_ polypeptide. (b) HPLC of TRX-(PQQPY)_8_ solubilized in 70% formic acid (5 mg.mL^−1^) (left) and the acid reaction products purified through a C18-T cartridge (right).

**S3 Figure. (a)** Association: two or more molecules bind to each other. Steady state: the number of molecules that is binding is equal to the amount of bonds that is breaking. Dissociation: the breaking of the bounds between the molecules. (b) The kinetic parameters (k_a_: association rate; k_d_: dissociation rate; K_D_=k_d_/k_a_: equilibrium dissociation constant) were determined by globally fitting the experimental data using BIA-evaluation software 4.1 (GE Healthcare) with available binding models (1:1 Langmuir model). The binding constants were calculated by two methods: affinity analysis (steady state fit) and association/dissociation analysis (kinetic fit).

**S4 Figure. Control of post-translational processing in transgenic lines.** (a): RP-HPLC of PINA from transgenic line A4 overexpressing PINA (red line) and from null segregant A5NS (blue line).(b) ratio of the integrated peak surfaces corresponding to the two major post-translational processed PINA, PINA1 and PINA2. Mass spectrometry characterization of the different peaks was previously described [20].

